# Rapid motor adaptation to bounce perturbations in online Pong game is independent from the visual tilt of the bouncing surface

**DOI:** 10.1101/2022.02.16.480739

**Authors:** Laura Mikula, Bernard Marius ’t Hart, Denise Y. P. Henriques

## Abstract

Motor adaptation describes the ability of the motor system to counteract repeated perturbations in order to reduce movement errors. Most research in the field investigated adaptation in response to perturbations affecting the moving hand. Fewer studies looked at the effect of a perturbation applied to the movement target, however they used simplistic visual stimuli. In this study, we examined motor adaptation to perturbations affecting the motion of dynamic targets. In addition, we asked whether external visual cues in the environment could facilitate this process. To do so, participants were asked to play an online version of the Pong game in which they intercepted a ball bouncing off a wall using a paddle. A perturbation was applied to alter the post-bounce trajectory of the ball and the wall orientation was manipulated to be consistent or not with the ball trajectory. The “trained tilt” group (n = 34) adapted to the consistent condition and the “trained horizontal” group (n = 36) adapted to the inconsistent condition. In case participants optimally integrate external visual cues, the “trained tilt” group is expected to exhibit faster and/or more complete adaptation than the “trained horizontal” group. We found that the perturbation reduced interception accuracy. Participants showed large interception errors when the perturbation was introduced, followed by rapid error decrease and aftereffects (errors in the opposite direction) once the perturbation was removed. Although both experimental groups showed these typical markers of motor adaptation, we did not find differences in interception success rates or errors between the “trained tilt” and “trained horizontal” groups. Our results demonstrate that participants quickly adapted to the dynamics of the pong ball. However, the visual tilt of the bouncing surface did not enhance their performance. The present study highlights the ability of the motor system to adapt to external perturbations applied to a moving target in a more dynamical environment and in online settings. These findings underline the prospects of further research on sensorimotor adaptation to unexpected changes in the environment using more naturalistic and complex real-world or virtual reality tasks as well as gamified paradigms.

## Introduction

Our motor system is incredibly efficient at generating precise motor actions, yet it also must retain some flexibility to adapt to various changing conditions. Motor adaptation refers to gradual adjustments of motor behavior in response to changes in task requirements or perturbations in the environment (Martin et al., 1996). During this process, the brain uses error signals in order to improve the accuracy of subsequent movements (Shadmehr et al., 2010; Wolpert et al., 2011).

Some of the most commonly used paradigms to study motor adaptation are force field and visuomotor rotation tasks. In force field adaptation, participants reach towards targets while their hand trajectory is deviated from the intended path by a robotic device that applies perturbing forces to the arm (Lackner & Dizio, 1994; Shadmehr & Mussa-Ivaldi, 1994). During visuomotor rotation, the cursor representing the visual feedback of the hand is rotated as participants are reaching to targets (Cunningham, 1989; Krakauer et al., 2000). Hence, adaptation to both force field and visuomotor rotation is driven by errors resulting from perturbations applied to the moving hand. In real life however, it seems more likely to face changes coming from the external environment rather than from our own body and effectors.

Interestingly, other types of experiments have allowed researchers to investigate how the motor system responds to perturbations affecting the target rather than the motor effector. Double-step, or “target jump”, tasks were initially used to study saccadic adaptation but have also been transposed to arm movements (Day & Lyon, 2000; Goodale et al., 1986). In double-step paradigms, participants are presented with a visual target which is displaced to another location at or after movement onset. Previous studies showed that, in response to target jumps, participants progressively reduced reach errors and showed aftereffects (i.e., reach errors in the opposite direction) once the perturbation was removed (Cameron et al., 2010; Laurent et al., 2011; Magescas & Prablanc, 2006; Westendorff et al., 2015). These findings demonstrate that we can adapt to visual perturbations applied to objects in our environment.

Nevertheless, these sensorimotor adaptation studies are often conducted in highly controlled laboratory settings using simplistic, isolated, and mostly static stimuli. In more naturalistic conditions, we would expect dynamic moving targets and possibly additional external cues to help infer their movement. One might then wonder what motor adaptation looks like when a perturbation is applied to a moving object that we interact with. More specifically, does the nervous system take into consideration visual cues in the surrounding environment to reduce movement errors and correct subsequent motor commands?

To investigate this question, we used an online version of the Pong game in which participants had to intercept a bouncing ball using a paddle controlled by their cursor. The Pong task is easy to implement and straightforward for participants to perform. Moreover, it has previously been used as a tool to study sensorimotor adaptation by altering the mapping between the hand and the paddle displayed on the screen. This was done by either introducing a delay (Avraham et al., 2017, 2019) or applying a rotation to the paddle relative to the hand position (Reichenthal et al., 2016). However, to our knowledge no studies have yet looked at the effect of a visual perturbation applied to the target (i.e., the moving ball) on the control of hand movements while playing Pong.

In the present study, the path of the pong ball was modified after it bounced off the upper wall (i.e., bouncing wall) so that participants would miss it. In addition, we manipulated the orientation of the bouncing wall to be congruent or not with the post-bounce ball trajectory. If participants effectively use visual cues of the surrounding environment, we should expect faster and/or greater sensorimotor adaptation when the tilt of the bouncing wall is consistent with the ball trajectory, as opposed to when the wall remains horizontal.

## Methods

### Participants

In total, 75 participants completed the experiment but 5 of them were excluded due to inconsistencies in the timing of stimulus presentation. For those 5 participants, the pong ball reached its bounce location more than 100 ms earlier (or later) than the median time, which might have made the task more (or less) difficult for them. For the other participants, the within-subject variability in bounce timings was on average 14 ms which is faster than the duration of one frame at 60 frames per second. Therefore, data of 70 participants was kept for analyses (mean age ± SD = 21.0 ± 4.3, range = 17–40; 12 males, 55 females, 2 identified as other, and 1 preferred not to say). Sixty-six participants self-identified as right-handed, 2 as left-handed, and 2 as ambidextrous. All participants reported having normal vision or being able to see their screen clearly.

Participants were university students recruited through the Undergraduate Research Participant Pool at York University, and they received credits as compensation for their participation. All of them gave informed electronic consent prior to participating. All procedures were in accordance with institutional and international guidelines, and were approved by York University’s Human Participants Review Committee.

### Apparatus

Participants accessed the study through the online survey platform Qualtrics (https://www.qualtrics.com) and they first answered questions about their demographics, health, and lifestyle. Then, participants were directed to Pavlovia where the task was hosted (demo version here: https://run.pavlovia.org/smcl/DemoPongTask). The experiment was created using PsychoPy3 Experiment Builder version 2021.1.4 (Peirce et al., 2019). PsychoPy has been shown to achieve good timing precision for visual stimulus presentation in online studies using different browser/operating system combinations (Bridges et al., 2020). Participants used their own computer to do the experiment. To move their cursor, 48 participants used a trackpad, 21 used a computer mouse, and one participant used a touchscreen. The display refresh rate was set to 60 Hz as it is the standard for most laptop and desktop monitors. The experiment was run in a web browser window in full screen mode.

The positions of visual stimuli were defined in a Cartesian coordinate system with the origin (0,0) at the center of the screen. Negative values represent down/left while positive values represent up/right. To accommodate for different screen resolutions, we used “height units” in PsychoPy (hereafter called arbitrary units a.u.) that scale visual stimuli relative to the height of the screen. In this system, the upper and lower edges of the screen always correspond to +0.5 and −0.5 a.u., respectively. Since the position of the left and right edges of the screen would vary depending on the aspect ratio of the monitor (e.g., ±0.8 for 16:10), we also scaled the cursor movements in the horizontal axis so that the far most left and the far most right positions correspond to −0.5 and +0.5 a.u., respectively. Hence, the experimental task was displayed in the central portion of the screen (size 1×1 a.u.).

### Pong task

Participants had to intercept a ball bouncing off a wall using a paddle. They used their cursor to move the paddle horizontally across the screen while the paddle’s vertical position remained unchanged. The goal of participants was to earn as many points as possible: They were instructed that they will get the most points if they intercept the ball using the central part of the paddle. Participants were free to use their preferred hand to complete the task and no viewing distance was imposed.

To start each trial, participants positioned their paddle (length = 0.1 a.u.) on one side of the screen, which was indicated by an arrow. Half of the trials started from the left and the other half from the right. Then, the pong ball (diameter = 0.025 a.u.) appeared at the same height and on the same side of the screen as the paddle. From this start position the ball moved upwards at a launching angle of 70°, 75°, 80°, or 85° to the horizontal. The X starting position of the ball was calculated so that the ball would always contact the horizontal wall in its center. Afterwards, the ball bounced back off the wall downwards and participants had to catch the ball with the paddle before it moved out of bounds. Participants got 10 points if the ball contacted the middle half of the paddle (0.05 a.u. = twice the size of the ball), 5 points if the ball contacted the paddle elsewhere, and 0 points if they missed the ball (see Figure 1A). At the end of each trial, participants received feedback on their performance and a score counter was displayed on the screen.

**Figure 1:**
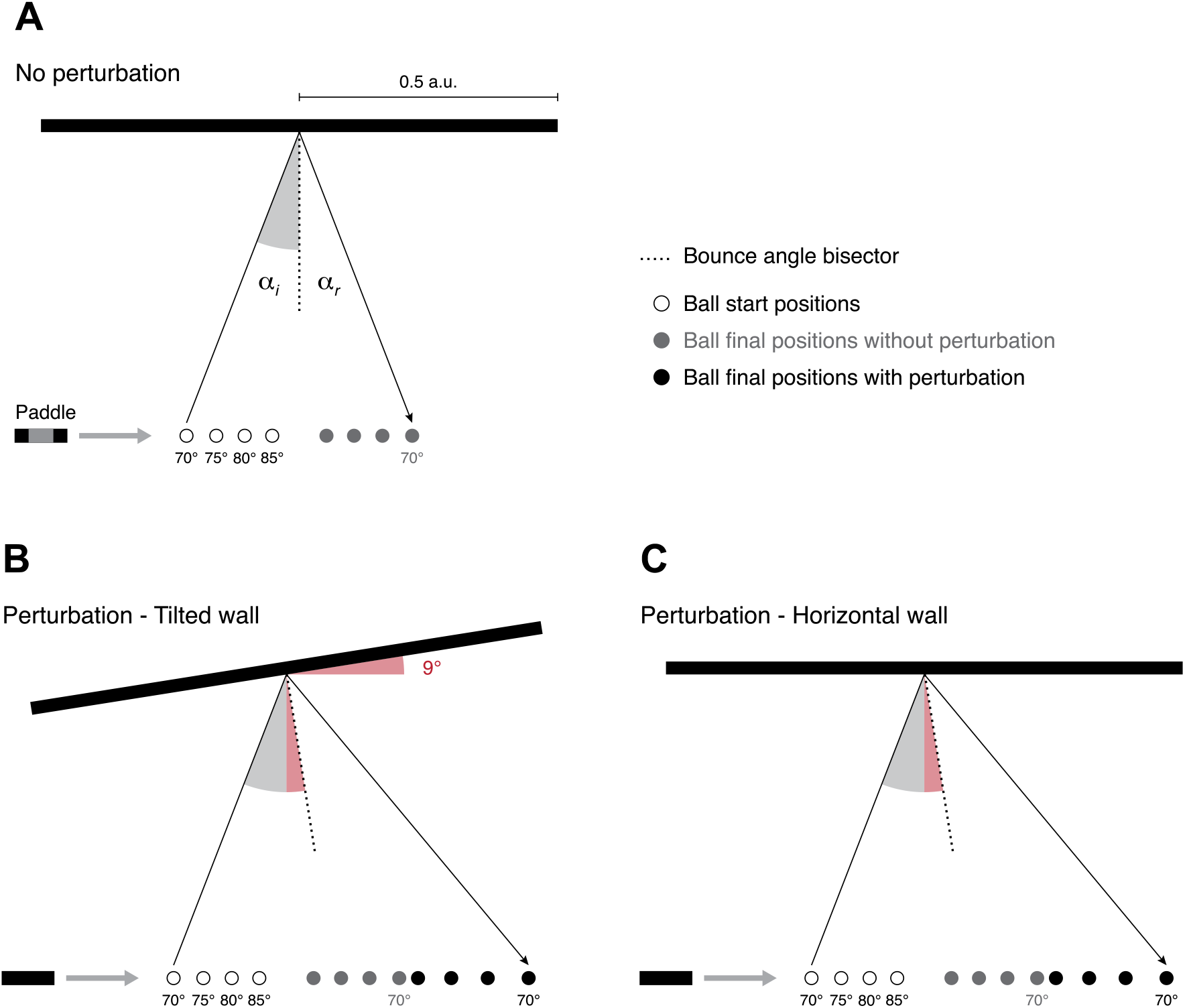
Experimental conditions. **A. Trials without perturbation.** The bounce angle bisector is aligned with the perpendicular to the wall and the ball final positions are the mirror of the start positions. During interception, participants get the maximum points if the ball contacts the grey area of the paddle (not visible to participants). **B. Trials with perturbation and tilted wall.** The wall is tilted by 9° and the bounce angle bisector stays aligned with the perpendicular to the wall. The ball final positions are further away from the paddle, but they are consistent with the wall orientation. **C. Trials with perturbation and horizontal wall.** Everything is the same as in the previous condition except that the wall is horizontal. Thus, the ball final positions are no longer consistent with the orientation of the wall. Dimensions in all figures are to scale.

### Experimental procedure

The bounce angle of the ball on the wall is the sum of the incident angle (between the upcoming ball and the perpendicular to the wall) and the reflected angle (between the departing ball and the perpendicular to the wall). Under the “no perturbation” condition, the reflected angle was equal to the incident angle. Therefore, the bounce angle bisector aligned with the perpendicular to the wall and the ball final positions were the mirror image of the ball start positions (Figure 1A).

In the “perturbation” condition, an angle of 9° was added to both the incident and reflected angles. This manipulation modified the ball trajectory only after contacting the wall. As a result, the final positions of the ball were displaced laterally by 0.25, 0.23, 0.21, and 0.20 a.u. for the 70°, 75°, 80°, and 85° launching angles respectively. Along with the perturbation, the orientation of the wall could be either tilted or stay horizontal. When the wall was tilted, the bounce angle bisector was aligned with the perpendicular to the wall. Therefore, the orientation of the wall was consistent with the trajectory of the ball (Figure 1B). If the wall was horizontal, that same ball trajectory became inconsistent with the visual scene, which should make the task more difficult for participants (Figure 1C).

The experiment was divided into two sessions separated by at least 12 hours. During the first session, participants completed 4 blocks of trials without perturbation followed by 4 blocks of trials with perturbation. Participants in the “trained horizontal” group (n = 36, 21.3 ± 4.4 years old) saw the horizontal wall during perturbation blocks while participants in the “trained tilt” group (n = 34, 20.7 ± 4.3 years old) saw the tilted wall (Figure 2).

**Figure 2:**
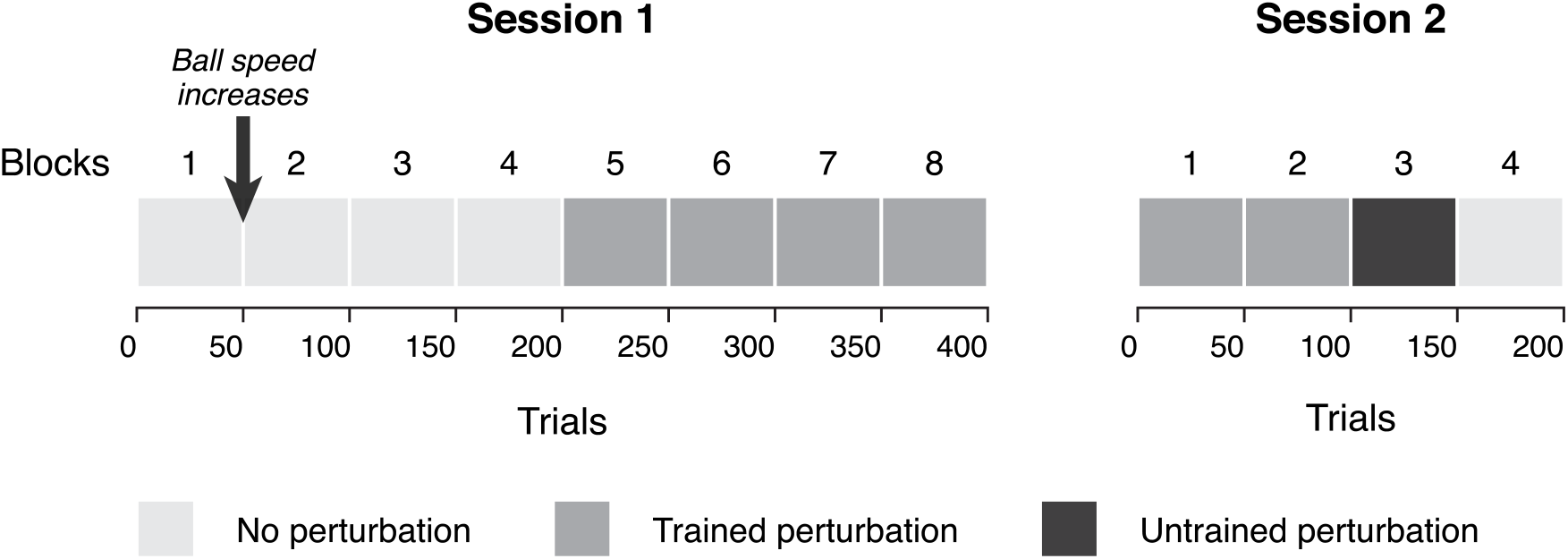
Experimental protocol. Session 1 consists of 4 blocks without perturbation and 4 blocks of trained perturbation (horizontal wall for the “trained horizontal” group and tilted wall for the “trained tilt” group). Session 2 includes 2 blocks of trained perturbation, followed by 1 block of untrained perturbation (tilted wall for the “trained horizontal” group and horizontal wall for the “trained tilt” group) and 1 block without perturbation. Each block comprises 50 trials.

The second session started with 2 blocks of the trained perturbation (the same perturbation condition as in the previous session). During block 3, we presented participants with the untrained perturbation condition: the “trained horizontal” group saw the tilted wall whereas the “trained tilt” group saw the horizontal wall. In block 4, the perturbation was removed for both groups and the orientation of the wall was horizontal. There were overall fewer participants (n = 46) who completed the second session (“trained horizontal” group: n = 25, 20.6 ± 3.8 years old; “trained tilt” group: n = 21, 20.5 ± 5.0 years old).

Participants were never informed that a perturbation was applied to the pong ball trajectory. Each block was composed of 50 trials in which the ball starting positions (left or right) and launch angles (70°, 75°, 80°, or 85°) were intermixed. The speed of the ball was set so that the time of arrival at the final location was constant, irrespective of the launching angle and the perturbation. During the first block of session 1, the pong ball reached the bounce location in 548 ms and the final location 567 ms later. In all the other blocks, the ball arrived at the bounce location in 399 ms and reached the final location 417 ms later.

### Data analyses

The interception success rate was computed for each participant as the proportion of trials in which the paddle contacted the ball with respect to the total number of trials in each block. The differences between conditions were assessed using two-way mixed ANOVA with group (“trained horizontal” or “trained tilt”) as a between-subject factor and block order as a within-subject factor. Bonferroni-corrected planned contrasts were used to compare interception success rates between specific blocks. During session 1, we specifically looked at the last block without perturbation (block 4), the first block of trained perturbation (block 5), and the last block of trained perturbation (block 8). During session 2, we were interested in the last block of trained perturbation (block 2), the block of untrained perturbation (block 3), and the block with no perturbation (block 4).

For each trial, the interception error was calculated as the difference between the ball final position and the center of the paddle, at the time the ball crossed the paddle plane. Interception errors were expressed in a.u., negative values indicate that the paddle did not move far enough (i.e., the position of the ball was underestimated) and positive values indicate that the paddle moved too far (i.e., the position of the ball was overestimated). For statistical analyses we only considered a subset of trials in each session. In session 1, we looked at the last trial without perturbation (trial 200), the first trial of trained perturbation (trial 201), and the 50th trial of trained perturbation (trial 250). In session 2, we were interested in the first and last trials of trained perturbation (trials 1 and 100), untrained perturbation (trials 101 and 150), and without perturbation (trials 151 and 200). The differences between conditions were assessed using two-way mixed ANOVA with group as a between-subject factor and trial as a within-subject factor. Bonferroni-corrected planned contrasts were used for follow-up post-hoc comparisons.

For each participant, trials in which the pong ball reached its bounce location more than 100 ms earlier or later than the median time were excluded (0.22% of trials removed). Trials in which the absolute value of interception errors was greater than 0.5 were excluded as well (1.10% of trials removed). Those were trials in which the paddle did not move or started to move very late relative to trial start. Data processing and statistical analyses were performed in R version 4.0.3 (R Core Team, 2020). For all statistical tests, the alpha level was set to 0.05. Departures from sphericity were adjusted using Greenhouse-Geisser corrections. Effect sizes for the ANOVAs are reported as partial eta-squared (*η*^2^_p_).

## Results

### Interception success rates

The success rates of participants during session 1 is depicted in Figure 3A. Note that the small dip in performance observed on block 2 is explained by the ball speed being faster than during block 1 (see the Methods section). Statistically there was no significant difference between the “trained horizontal” and the “trained tilt” groups (*F*_(1,68)_ = 2.37, *p* = 0.129), and no interaction between block order and group (*F*_(7,476)_ = 2.08, *p* = 0.052). However, we found a significant effect of block order (*F*_(7,476)_ = 136.24, *p* < 0.001, *η*^2^_p_ = 0.667). As shown by post-hoc comparisons, the overall success rate of participants was 76.4% during the last block without perturbation and it decreased when the trained perturbation was first introduced (53.4%; *t*_(68)_ = 16.05, *p* < 0.001, *d* = 2.97). By the last block of trained perturbation, the interception success rate increased to 60.8% (*t*_(68)_ = 11.89, *p* < 0.001, *d* = 2.02) but still remained lower than without perturbation (*t*_(68)_ = 6.10, *p* < 0.001, *d* = 0.95). This demonstrates that, irrespective of the groups, the bounce ball perturbation decreased interception performance. Furthermore, although participants improved over time, their performance did not return to baseline levels by the end of session 1.

**Figure 3A and 3B:**
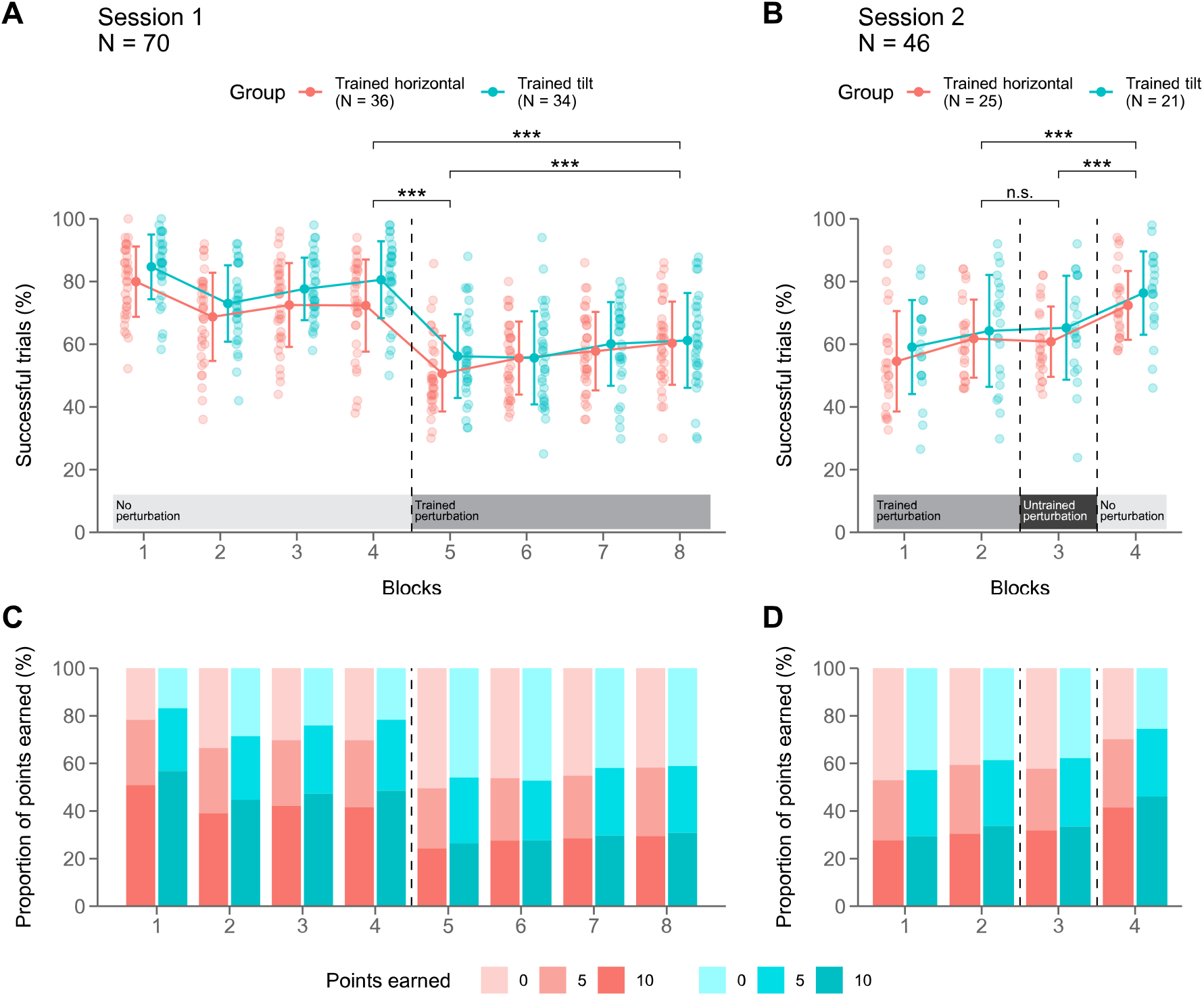
Interception success rates. Proportion of trials in which participants intercepted the ball. Error bars correspond to ± 1 SD. \*\*\**p* < 0.001, n.s.: non-significant. **3C and 3D: Proportions of points earned.** Participants got 10 points when they intercepted the ball using the middle half of the paddle, 5 points when using the sides of the paddle, and 0 points if they missed the ball.

In session 2 (Figure 3B), we found neither an effect of group (*F*_(1,44)_ = 1.07, *p* = 0.307), nor an interaction between block order and group (*F*_(3,132)_ = 0.18, *p* = 0.898). In contrast, there was a significant effect of block order on interception success rates (*F*_(3,132)_ = 41.11, *p* < 0.001, *η*^2^_p_ = 0.483). More specifically, success rates did not change when participants switched from the trained perturbation (63.0%) to the untrained perturbation (63.0%; *t*_(44)_ = 0.002, *p* = 1.00). Once the perturbation was removed, the interception success rate increased to 74.3%, which was significantly higher than success rates in both the untrained perturbation block (*t*_(44)_ = 8.04, *p* < 0.001, *d* = 1.47) and the second block of trained perturbation (*t*_(44)_ = 8.11, *p* < 0.001, *d* = 1.47). These results suggest that switching perturbations between both groups did not affect interception performance. However, participants’ performance improved when the bounce ball perturbation was removed.

### Proportion of points earned

The proportions of points earned by participants in each block are shown in Figures 3C and 3D. Participants earned 10 points when intercepting the ball using the middle half of the paddle, 5 points when using the sides of the paddle, and no points when they missed the ball. In session 1, Figure 3C shows that after the introduction of the perturbation (blocks 5 to 8), the decrease in overall success rate was associated with less trials in which the ball contacted the middle part of the paddle (10-point trials). Whereas the proportion of trials in which the ball contacted the sides of the paddle (5-point trials) stayed constant throughout the blocks. This was true for both the “trained horizontal” and the “trained tilt” groups. During session 2 (Figure 3D), the proportion of 10-point trials was similar in the trained and untrained perturbation conditions (blocks 1 to 3) and slightly increased when the perturbation was removed (block 4). In contrast, the proportion of 5-point trials overall remained the same across the four blocks. These observations suggest that the ball perturbation mainly affected interception accuracy as participants were less able to bring the center of the paddle to the ball final position.

### Interception errors

Average interception errors on each individual trial are depicted in Figure 4. During the first session (Figure 4A), participants in both the “trained horizontal” and “trained tilt” groups initially exhibited small interception errors on the very first few trials of block 1. As the no perturbation trials progressed, average errors became smaller than the length of the paddle. On the first trial of trained perturbation, participants showed large interception errors about half the size of the perturbation displacement. The errors were negative, indicating that participants did not move their paddle far enough to intercept the moving ball. Then, interception errors reduced within 10 to 15 trials and remained stable until the end of the session. When conducting a mixed ANOVA with group and trial (trials 200, 201, and 250) as factors, we found a significant effect of trial (*F*_(2,132)_ = 43.13, *p* < 0.001, *η*^2^_p_ = 0.395) but no significant effect of group (*F*_(1,66)_ = 0.78, *p* = 0.379), and no interaction between group and trial (*F*_(2,132)_ = 0.14, *p* = 0.857).

**Figure 4A and 4B:**
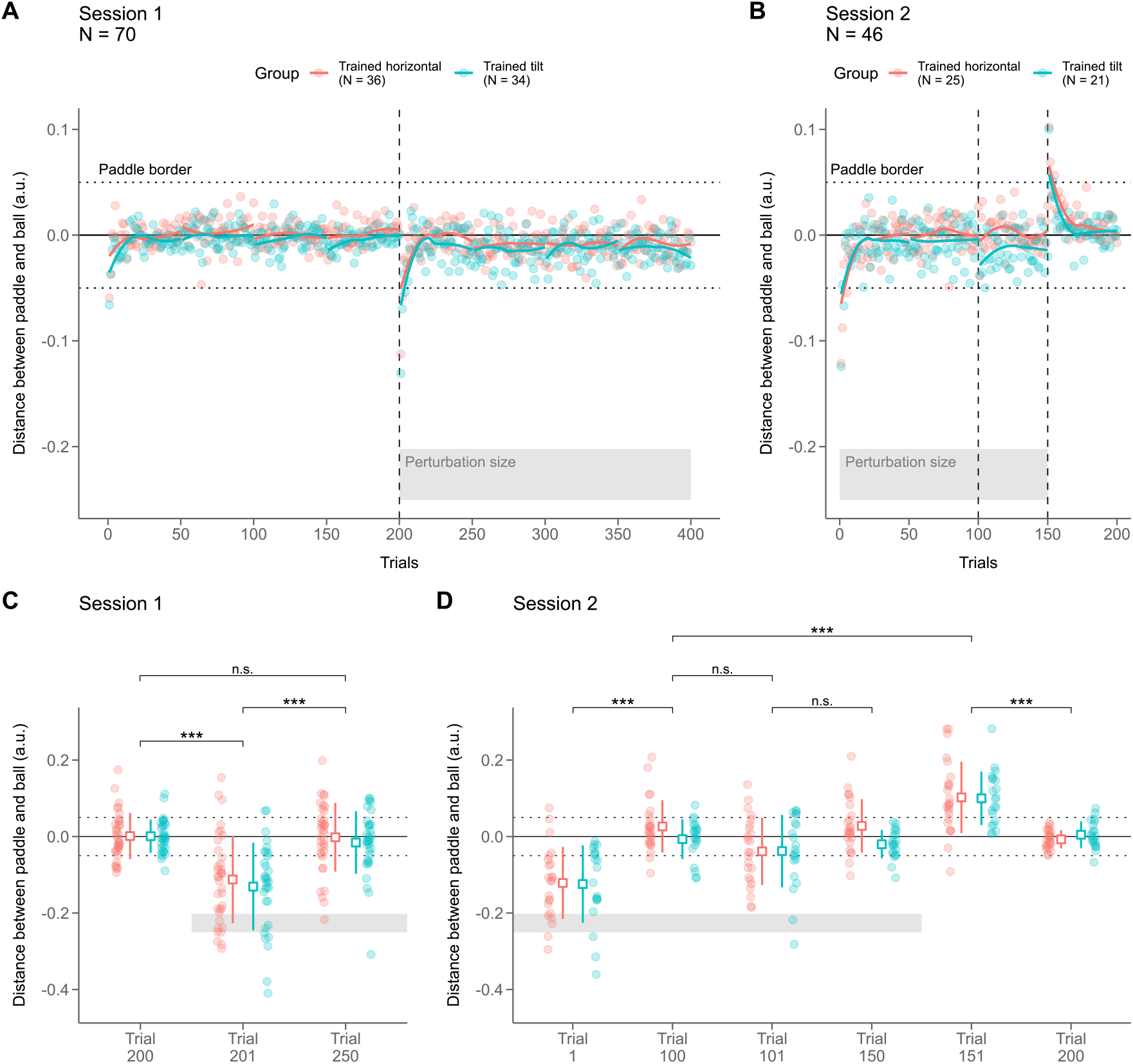
Interception errors across individual trials. Distance (in a.u.) between the center of the paddle and the final position of the ball. Values are negative when participants did not move the paddle far enough and positive when they moved the paddle too far. Circles correspond to the interception errors averaged across participants. The colored lines are the regression lines fitted to each group and each block of 50 trials. Dotted lines depict the length of the paddle, and the grey area represents the size of the perturbation. **4C and 4D: Statistical analyses.** Interception errors during trials of interest. White squares represent the mean interception error across participants in each group and error bars correspond to ± 1 SD. \*\*\**p* < 0.001, n.s.: non-significant.

Post-hoc comparisons (Figure 4C) showed a significant increase in interception error between the last trial with no perturbation and the first trial in which the perturbation was introduced (trials 200 vs. 201; *t*_(66)_ = 7.76, *p* < 0.001, *d* = 1.43). By the end of the first block of trained perturbation, errors were significantly reduced (trials 201 vs. 250; *t*_(66)_ = 7.45, *p* < 0.001, *d* = 1.33) and were not different from interception errors in the last trial without perturbation (trials 200 vs. 250; *t*_(66)_ = 0.63, *p* = 1.00).

During the second session (Figure 4B) participants in the “trained horizontal” and “trained tilt” groups showed large errors on the first trial of trained perturbation followed by a quick error reduction, similar to what we observed in session 1. Switching from the trained to the untrained perturbation did not seem to affect interception errors, irrespective of the group. Finally, participants exhibited large positive errors when the perturbation was removed, indicating that they moved the paddle too far relative to the ball final position. These errors in the opposite direction than initial errors are known as aftereffects and are characteristic of motor adaptation. Aftereffects then quickly decayed with a rate similar to that of the initial adaptation errors. A mixed ANOVA with group and trial (trials 1, 100, 101, 150, 151, and 200) as factors showed a significant effect of trial (*F*_(5,200)_ = 41.66, *p* < 0.001, *η*^2^_p_ = 0.510) but no significant effect of group (*F*_(1,40)_ = 2.03, *p* = 0.162), and no interaction between group and trial (*F*_(5,200)_ = 1.34, *p* = 0.261).

As shown by post-hoc comparisons (Figure 4D), interception errors in the first trial of trained perturbation were significantly reduced by the end of the second block of trained perturbation (trials 1 vs. 100; *t*_(66)_ = 7.24, *p* < 0.001, *d* = 1.77). These errors were not modulated by the introduction of the untrained perturbation (trials 100 vs. 101; *t*_(66)_ = 2.32, *p* = 0.127). There was no significant change in interception error between the beginning and the end of the untrained perturbation block (trials 101 vs. 150; *t*_(66)_ = 2.20, *p* = 0.168). Finally when the perturbation was removed, participants made positive interception errors that were larger than those observed after they adapted to the trained perturbation (trials 151 vs. 100; *t*_(66)_ = 6.06, *p* < 0.001, *d* = 1.33). The aftereffects then decreased close to 0 on the last trial without perturbation (trials 151 vs. 200; *t*_(66)_ = 6.89, *p* < 0.001, *d* = 1.38).

Additionally, we looked at savings which corresponds to faster adaptation when reexposed to a previous perturbation. Thus, we tested whether initial errors to the trained perturbation were smaller in session 2 than in session 1 (session 1 trial 210 vs. session 2 trial 1). The analysis was made using only participants who completed both sessions. The mixed ANOVA with group and session as factors showed no main effects of group (*F*_(1,40)_ = 0.13, *p* = 0.721) or session (*F*_(1,40)_ = 0.37, *p* = 0.545), and no interaction effect (*F*_(1,40)_ = 0.72, *p* = 0.400). Theses result suggest that there was no savings in neither of the experimental groups.

## Discussion

In the present study, participants played an online Pong game where they had to intercept a moving ball using a paddle controlled by their cursor. A fixed rotation was applied to the pong ball trajectory after it contacted the bouncing wall, and the tilt of the wall was modified to be consistent (tilted wall) or inconsistent (horizontal wall) with the post-bounce path of the ball. We hypothesized that, if visual cues in the surrounding environment are integrated by the nervous system, motor adaptation should be enhanced when the bouncing wall is tilted and congruent with the ball trajectory. To test this, we had two groups adapting to either the consistent (“trained tilt” group) or the inconsistent condition (“horizontal tilt” group) on a first session. During the subsequent session, we assessed motor adaptation savings, transfer when switching to the other (untrained) perturbation, as well as aftereffects when perturbations were removed.

Both the “trained horizontal” and “trained tilt” groups in our experiment showed clear markers of sensorimotor adaptation. When the pong ball perturbation was introduced, participants first made large errors which then gradually decreased with time. In addition, participants showed strong aftereffects (i.e., errors in the direction opposite to the disturbance) after the perturbation was removed (Kluzik et al., 2008; Lackner & Dizio, 1994; Martin et al., 1996). Savings on the other hand refers to faster sensorimotor adaptation, and sometimes smaller initial errors, following reexposure to a previously experienced perturbation. In this study, we did not observe savings between the first and the second sessions, which is in contrast with previous findings (Klassen et al., 2005; Krakauer et al., 2005). Our results nevertheless suggest some short-term retention (Figures 4A and 4B). Indeed, within the same session, there was no increase in initial interception errors between successive blocks of trained perturbation.

We found that the perturbation we applied to the ball trajectory affected mainly the ability of participants to intercept the ball with the central part of the paddle (i.e., higher interception accuracy). This observation is probably related to the fact that participants did not make ballistic movements and were able to make online corrections when moving the paddle using their cursor. That might explain the fast adaptation rate to the pong ball’s dynamics and how interception errors were reduced within 10 to 15 trials. In addition to online error corrections, a few studies have reported within-trial adaptation to visuomotor rotations (Braun et al., 2009) and force field (Crevecoeur et al., 2020). Because our measure of interception errors is related to the final position of the paddle, it is likely to reflect a combination of trial-by-trial adaptation, online feedback correction, and perhaps within-trial adaptation. However, we argue that such a continuous control process is closer to what happens in more ecological conditions. Lastly, since our experiment was not designed to dissociate implicit and explicit components of adaptation, we cannot rule out the possibility that participants relied on more cognitive strategies to complete the task. Nevertheless, the aftereffects present in this study indicate a role of implicit adaptation and rather suggest a combination of both implicit and explicit processes.

It has been proposed that sensorimotor adaptation is related to an internal forward model, generated by the nervous system, which predicts the sensory consequences of motor commands. In case of a perturbation, sensory prediction errors (mismatch between expected and actual sensory feedback) are used to update this internal model and reduce movement errors (Izawa & Shadmehr, 2011; Shadmehr & Mussa-Ivaldi, 1994; Tseng et al., 2007; Wolpert & Miall, 1996). It has first been argued that motor adaptation is mostly driven by sensory prediction errors whereas target errors (discrepancy between the target and the movement feedback) have little to no involvement in this process (Diedrichsen et al., 2005; Mazzoni & Krakauer, 2006; Taylor & Ivry, 2011). This view was challenged by several target jump studies demonstrating reach adaptation in response to visual target displacements (Cameron et al., 2011; Laurent et al., 2011; Magescas et al., 2009; Magescas & Prablanc, 2006). Though it was pointed out that, when noticed by participants, target jumps fail to induce adaptation and rather lead to re-aiming strategies (Cameron et al., 2010; Westendorff et al., 2015). Thus, methodological differences might explain apparently conflicting results obtained in previous studies discarding the role of target errors in adaptation.

The results from our study further support the evidence that adaptation of internal models is also sensitive to target errors (Reichenthal et al., 2016). As opposed to most previous studies with targets jumping from one location to another, we used continuously moving targets that participants had to intercept. In this particular condition, successful interceptions rely on the ability of internal models to predict the target’s dynamics. This claim is supported by a previous study investigating eye movements in a virtual-reality interception task (Diaz et al., 2013). The authors found that participants accurately predicted the position of the bouncing balls even though their post-bounce trajectories were altered by changes in ball elasticity. They concluded that the control of eye movements depends not only on currently available visual information but also on experience-based models of dynamical properties of the moving object. Our results suggest a similar process in which participants use an internal model of the dynamics of the bouncing ball that is updated based on prior knowledge, as it has already been proposed for the interception of objects falling under gravity (Zago et al., 2009).

Despite indications of adaptation, we found no evidence supporting our hypothesis that external visual cues enhance motor adaptation to perturbations of moving targets. Throughout the experiment, the two groups performed similarly; there was no significant difference in the speed or magnitude of adaptation. Also contrary to our predictions, the “trained tilt” group which first adapted to the consistent condition did not show any benefits when switching to the inconsistent condition and savings was not larger on the second session. Altogether these findings suggest that participants adapted to the dynamical properties of the pong ball, but that the visual tilt of the bouncing surface did not improve their performance. The control of interceptive actions seems to rely on an internal model of the dynamics of the target itself, irrespective of the surrounding environment. This could be explained by the visual information being processed through two major pathways: the ventral stream more involved in perception and the dorsal stream more involved in action (Goodale & Milner, 1992). For instance, participants have been asked to intercept a moving disc at its bounce location using a paddle. While the interceptive movements of participants were accurate, their perceptual judgments about the bounce location were consistently biased (Marinovic et al., 2012). Alternatively, the visual tilt of the wall may have been deemed irrelevant for this specific task. It would be interesting to see if different results are obtained when participants need to interact with the wall. For example, if they are asked to choose where to bounce the ball on the wall so that it reaches a particular location on the opposite side.

In conclusion, the results from this study show that sensorimotor adaptation to target errors is possible in an online Pong game. It supports the findings of previous studies demonstrating that online experimentation can be as informative if not more than traditional laboratory experiments (Kim et al., 2021; Tsay et al., 2021). The Pong task is less restrictive than the ones that have been used in past lab-based studies. Participants are able to move more freely (although in our version, motion was restricted to one dimension) and they have to figure out where and when to move to achieve their goal. Our findings encourage for further investigation on sensorimotor adaptation in more naturalistic and dynamic environments, as well as the use of more gamified tasks for research purposes.

